# Fabrication of a Parali (Rice Straws) Biomass & Herbal Based Filtration System for Sustainable Rural Water Purification

**DOI:** 10.1101/2025.10.13.682254

**Authors:** Kishlay Kant Singh, Divya Prakash, Mansi Saini

## Abstract

Access to safe drinking water remains a serious challenge in rural India, where traditional chemical and membrane-based filtering systems are often unavailable due to their high cost, energy requirements, and maintenance demands. This work develops and validates a low-cost, eco-friendly herbal filtration system for Mubarakpur hamlet by combining the antibacterial properties of plant extracts with natural filter media. The physical, chemical, and microbiological properties of the water samples collected from ponds, wells, sewage, and tap sources included turbidity, pH, COD, BOD, heavy metals, and bacterial load. Herbal extracts derived from Neem (Azadirachta indica), Bael (Aegle marmelos), Lemon, Amla, Moringa, Garlic, and Ginger had bactericidal effects when tested for antimicrobial activity; however, the peels of Bael and Lemon exhibited the best inhibition of Streptobacillus species. A six-layer prototype filtration device including cotton cloth, charcoal, parali biomass, gravel, sand, and a layer of herbal extract was created as a result of these findings. The extracts of lemon and bael exhibited the greatest inhibition of Streptobacillus species when tested for antibacterial activity. A six-layer prototype filtration device including cotton cloth, charcoal, parali biomass, gravel, sand, and a layer of herbal extract was created as a result of these findings. Laboratory testing of the small prototype revealed significant improvements in water quality: the microbial load dropped from 2,000,000 CFU/mL to 0 CFU/mL, the turbidity went from 300 NTU to 0.8 NTU, the COD decreased from 1000 mg/L to 9 mg/L, and the BOD decreased from 200 mg/L to 3.5 mg/L. Lead levels in tap water (0.02 mg/L) were also reduced to undetectable levels.

## 1. Introduction

Enter Everyone acknowledges that access to safe, clean drinking water is essential for growth, health, and societal advancement and is a fundamental human right. Despite decades of national and international efforts, millions of people in rural India continue to suffer from water pollution. The primary sources of drinking water in communities, surface water and groundwater, are increasingly under risk from untreated sewage, agricultural runoff, industrial effluents, and natural contaminants like viruses and heavy metals. The findings are alarming: regular outbreaks of waterborne diseases including cholera, typhoid, dysentery, and diarrhoea continue to endanger the health and productivity of rural people. According to the World Health Organisation (WHO), over two billion people globally consume water contaminated by human waste, and a significant chunk of this population resides in India, where rural regions are disproportionately affected. This highlights the pressing need for environmentally friendly and locally adaptable purification methods [1].

### 1.1. Rural Water Issues and Current Constraints

Water pollution in rural areas is a complex issue with physical, chemical, and biological components. Water is physically unfit for residential use because to its excessive turbidity, disagreeable taste, and offensive scent. Chemically, safety is further jeopardised by the presence of heavy metals like lead, arsenic, and mercury, nitrates, phosphates, and pesticides. Pathogens such as Escherichia coli, Streptobacillus species, and coliform bacteria are biologically dangerous and can cause major health problems. In rural areas, untreated sewage interacting with surface water continues to be a significant source of pollution. In contrast to metropolitan areas that have centralised water treatment systems, rural areas frequently rely on uncontrolled sources such ponds, small rivers, and bore wells, which forces households to drink tainted water and prolongs sickness and financial cycles [2].

To deal with these problems, a number of traditional purification methods have been used, including membrane filtering, alum coagulation, and chlorination. Nevertheless, a number of constraints limit their use in rural areas. Despite its effectiveness, chlorination changes the taste and smell of water and can produce dangerous byproducts like trihalomethanes. Clarity is increased by alum coagulation, however microbiological pollutants are not removed. High-efficiency membrane technologies like reverse osmosis (RO) and ultrafiltration are expensive, energy-intensive, and generate enormous amounts of reject water that need to be disposed of safely [3].

Furthermore, these systems require regular maintenance, professional operation, and replacement parts, conditions that are seldom satisfied in rural areas with low resources. Long-term sustainability issues continue to exist notwithstanding technical constraints. Ecological and economic equilibrium are threatened by reliance on non-renewable resources and chemical chemicals. Thus, there is a growing need to investigate affordable, environmentally responsible, and locally applicable purification technologies that can successfully close the gap between rural accessibility and urban water safety regulations [4].

### 1.2. Herbal and natural materials: a promising alternative

Traditional knowledge and ethnobotanical techniques have long emphasised the antibacterial and medicinal properties of plants such as Amla (Phyllanthus emblica), Bael (Aegle marmelos), Neem (Azadirachta indica), and Moringa (Moringa oleifera). Neem in particular is widely used in Ayurvedic medicine because to its strong antibacterial, antiviral, and antifungal properties. Similarly, Bael leaves and fruit extracts are rich in bioactive compounds that neutralise toxins and stop the growth of microorganisms. Recent experimental studies have shown that plant-based extracts can effectively reduce turbidity and decrease waterborne pathogens without leaving behind dangerous residues [5].

These promising results notwithstanding, most research remains isolated, fragmented, and laboratory-scale. The antibacterial properties of specific plants have been well documented by pharmacological and microbiological research, but there is no scientific backing for integrating them into a workable, scalable water filtering system. The lack of standardised processes, thorough testing, and prototype development has hindered the transformation of herbal-based concepts into practical solutions for rural communities. This gap in the literature highlights the need for comprehensive research that not only demonstrates the efficacy of herbal extracts but also incorporates them into technology solutions suitable for real-world applications [6].

When taken as a whole, the evidence shows that conventional chemical and membrane-based approaches are inadequate for solving the persistent environmental and health issue of rural water pollution. Herbal-based purification has not yet undergone extensive testing or been integrated into engineering prototypes, despite its promise. This book bridges that gap by offering the development, validation, and analysis of a multi-layer herbal filtration system that combines plant extracts such as neem, bael, and others with agricultural wastes like charcoal and parali biomass [7]. By investigating water quality improvements, antimicrobial efficacy, and prototype performance, the project contributes to scientific understanding and offers practical pathways for sustainable, community-driven rural water purification [8].

## 2. Materials and Methods

In This work employed a multidisciplinary strategy that includes field surveys, laboratory analysis, the production of herbal extracts, and prototype fabrication in order to evaluate an inexpensive herbal-based filtration system for rural water purification. The study was carried out in the Indian village of Mubarakpur, where a lack of centralised treatment equipment and untreated sewage are the main causes of poor water quality [9]. Systematic sampling from ponds, wells, boreholes, sewage drains, and taps provided water for physicochemical, chemical, and microbiological analysis, and a structured household survey documented water-use patterns and community sentiments. As a consequence of these findings, herbal extracts were tested for antibacterial activity and added to a multi-layer prototype, which was then confirmed in a laboratory [10].

### 2.1. Study Area and Context

The study was conducted in the rural Indian hamlet of Mubarakpur, where the majority of the population gets their drinking water from boreholes, ponds, wells, and municipal taps. Preliminary surveys revealed widespread concerns with turbidity, odour, and the frequency of waterborne diseases. Importantly, the town was an ideal test site for evaluating a low-cost herb-based filtration system because it lacked a central water treatment plant [11].

### 2.2. Survey and Sampling

To establish a baseline, a comprehensive home survey was conducted between October and January. As part of the study, villagers were interviewed to learn their opinions on the water’s quality, the sources they used, and the frequency of waterborne infections. Following the survey, water samples were collected from a variety of sources, including tap water, ponds, sewage drains, boreholes, and wells. The sample, which was processed in 12 hours and delivered in refrigerators, was taken in sterile glass bottles [12].

### 2.3. Physical parameters

For every water sample, physical characteristics such as temperature, colour, turbidity, electrical conductivity, salinity, total dissolved solids (TDS), odour, taste, specific gravity, and transparency were assessed. A digital turbidimeter was used to quantify turbidity in NTU, and portable conductivity meters were utilised to assess TDS and electrical conductivity. To guarantee precise water quality characterisation, other criteria including salinity, odour, taste, and transparency were evaluated using accepted practices [13].

### 2.4. Chemical parameters

Chemical pH, dissolved oxygen (DO), COD, BOD, nitrates, nitrites, ammonia, phosphates, chlorine, hardness, and heavy metals were among the chemical parameters that were examined. A calibrated pH meter was used to test pH, the modified Winkler method was used to quantify BOD, and the dichromate reflux method was used to determine COD [14]. Heavy metals (Pb, As, Hg, and Cd) were identified using Atomic Absorption Spectroscopy (AAS), whereas nutrients such phosphates, ammonia, nitrates, and nitrites were measured spectrophotometrically [15].

### 2.5. Biological parameters

Using standard culture-based methods, microbial analysis was conducted. Colony-forming units (CFU/mL) were measured by plating water samples on selective medium and incubating them for 24 to 48 hours at 37 °C [16]. Biochemical assays such as catalase, oxidase, MR-VP, and indole tests were used in conjunction with Gramme staining to identify the bacteria. Enzyme activity, total microbial load, bacterial count, algal biomass, and nutrient content (phosphorus and nitrogen) were among the biological characteristics evaluated [17].

### 2.6. Herbal Extract Preparation

Based on literature and tradition, seven plants were selected for antimicrobial testing: Neem (Azadirachta indica), Bael (Aegle marmelos), Lemon (Citrus limon) peels, Amla (Phyllanthus emblica), Moringa (Moringa oleifera) leaves, Garlic (Allium sativum) peels, and Ginger (Zingiber officinale) peels [18]. Fresh plant materials growing nearby were collected, thoroughly cleaned, allowed to dry in the shade, and then milled into a powder. To extract it, 10 g of powdered material was soaked in 100 mL of distilled water for 24 and 48 hours. The extracts were filtered through muslin cloth and centrifuged to produce a clear supernatant [19].

#### 2.6.1. Antimicrobial testing & Prototype Development

The agar well diffusion method was used to evaluate the antibacterial activity against Streptobacillus isolates that were isolated from polluted water. To assess the effectiveness of several plant-based extracts, zones of inhibition were evaluated during a 24-hour incubation period at 37°C. In contrast to extracts soaked for just 24 hours, which demonstrated little antimicrobial activity, extracts soaked for 48 hours had considerably wider inhibitory zones. Amla and neem leaves showed significant bactericidal qualities after incubation, whereas bael leaves showed the highest inhibitory potential among the botanicals examined, followed by lemon peels. On the other hand, moderate inhibitory zones were created by moringa, ginger, and garlic, indicating that they serve as supplemental antimicrobial agents rather than principal inhibitors [20].

On the basis of survey data, chemical analysis, and antimicrobial efficacy testing, a six-layer prototype filter unit was created. A 1:4 scaled version of the proposed design was represented by the laboratory-scale model, which was built using a stainless steel mesh cuboidal frame that was 1 foot 4 inches long, 5 inches wide, and 2 inches high. Each of the six successive levels that made up the filtering system had a specific function in the purification of water [21]. The removal of coarse debris and suspended particles was made easier by the first gravel layer (5 cm). The purpose of the second layer of sand (5 cm) was to filter out tiny particles and reduce turbidity. The third layer served as a natural bio-absorbent for organic pollutants and was made up of agricultural leftovers known as Parali biomass (5 cm). The fourth layer, a herbal mixture (5 cm) made up of vegetable waste, citrus peels, Bael leaves, and Neem leaves, has antifungal and antibacterial properties. Heavy metals, chemical poisons, and odours were all removed by the fifth layer of charcoal (5 cm). Lastly, a 2 cm layer of cotton mesh strengthened the elimination of chemical impurities and odours and provided further filtering for any remaining pollutants [22].

## 3. Results & Discussion

Using sterilised glass bottles, water samples were taken from a variety of sources, such as ponds, sewage drains, boreholes, wells, and tap water. They were then chilled for processing within 12 hours. Water quality significantly improved after being treated with the herbal multi-layer prototype. These included a sharp drop in the microbial load from 2,000,000 CFU/mL to very low levels, as well as decreases in turbidity, chemical oxygen demand (COD), biological oxygen demand (BOD), and lead concentrations. The effectiveness of this inexpensive, environmentally friendly filtering method is further supported by the strong antibacterial activity of bael and lemon extracts. A thorough explanation of these results in relation to current literature and traditional methods is provided in the next subsections.

### 3.1. Baseline Water Quality Assessment

A detailed analysis of untreated water from ponds, wells, sewage drains, boreholes, and tap water revealed severe contamination. The sewage water’s 300 NTU turbidity, 1000 mg/L COD, and 200 mg/L BOD levels all indicated heavy organic pollution. Streptobacillus species predominated, according to microbiological testing, with about 2,000,000 CFU/mL. The amount of lead (0.02 mg/L) in tap water sparked worries about long-term harm. The combined risk of chemical and microbiological contamination in communities is highlighted in WHO/UNICEF publications on rural water pollution, which are consistent with our baseline findings [23].

### 3.2. Herbal Extract Antimicrobial Evaluation

Prior to filter integration, the antibacterial activity of aqueous extracts of seven botanicals; Bael, Neem, Lemon, Amla, Moringa, Garlic, and Ginger, was assessed against Streptobacillus moniliformis, which was found in significant quantities in the water sample. 48-hour-soaked extracts exhibited much larger inhibitory zones (22–28 mm) than 24-hour extracts (10–14 mm), suggesting that active phytochemicals diffused more readily. Neem had a definite bactericidal action, whereas bael leaves and lemon peels showed the highest inhibition. The overall synergistic antibacterial response was aided by the modest inhibition that amla, moringa, garlic, and ginger caused. This is the rating of potency: ***Amla > Moringa > Garlic > Ginger > Lemon > Neem > Bael***. Given that their combined effect allowed for total microbial eradication (0 CFU/mL) in treated water, these results (Table 2, Figures 7 & 8) support the justification for adding these extracts to the prototype and provide a sustainable and natural substitute for chemical disinfectants.

**Table 1:** Essential Chemical and Physical Parameter Results.

| *Parameter* | *Untreated Sewage Water* | *Treated Water* | *Permissible Limit (WHO / BIS Standard)* |
| --- | --- | --- | --- |
| Turbidity (NTU) | 300 | 0.8 | $\leq 1$ |
| COD (mg/L) | 1000 | 9 | $\leq 10$ |
| BOD (mg/L) | 200 | 3.5 | $\leq 5$ |
| Lead (mg/L) | 0.02 | Undetectable | $\leq 0.01$ |
| pH | 7.66 | 6.9 | 6.5 – 8.5 |
| Electrical Conductivity ($\mu\text{S}/\text{cm}$) | 2000 | 8 | $\leq 500$ |
| Color | Dark Brown/Black | Clear | Colorless |
| Odor & Taste | Strong, Unpleasant | Odorless/Tasteless | Acceptable / Odorless |
| Nitrate/Nitrite (mg/L) | 50 | $\leq 0.32$ | $\leq 45$ (Nitrate), $\leq 3$ (Nitrite) |
| Phosphate (mg/L) | 10 | 0.2 | $\leq 0.3$ |
| Hardness (mg/L) | 300 | 120 | $\leq 200$ (Desirable) |

**Table 2:** Herbal extracts of neem, moringa, ginger, garlic, bael, lemon, basil, and amla prevent the growth of multidrug-resistant (MDR) Streptobacillus moniliformis infection.

| *Herbal Material* | *Observed Activity* |
| --- | --- |
| 1. Bael Leaves | Strong inhibition (largest zone) |
| 2. Lemon Peels | Strong inhibition |
| 3. Neem Leaves | Bactericidal effect |
| 4. Amla Leaves & Fruits | Moderate bactericidal effect |
| 5. Moringa Leaves | Moderate inhibition |
| 6. Garlic Peels | Moderate inhibition |
| 7. Ginger Peels | Moderate inhibition |

**Figure 1:**
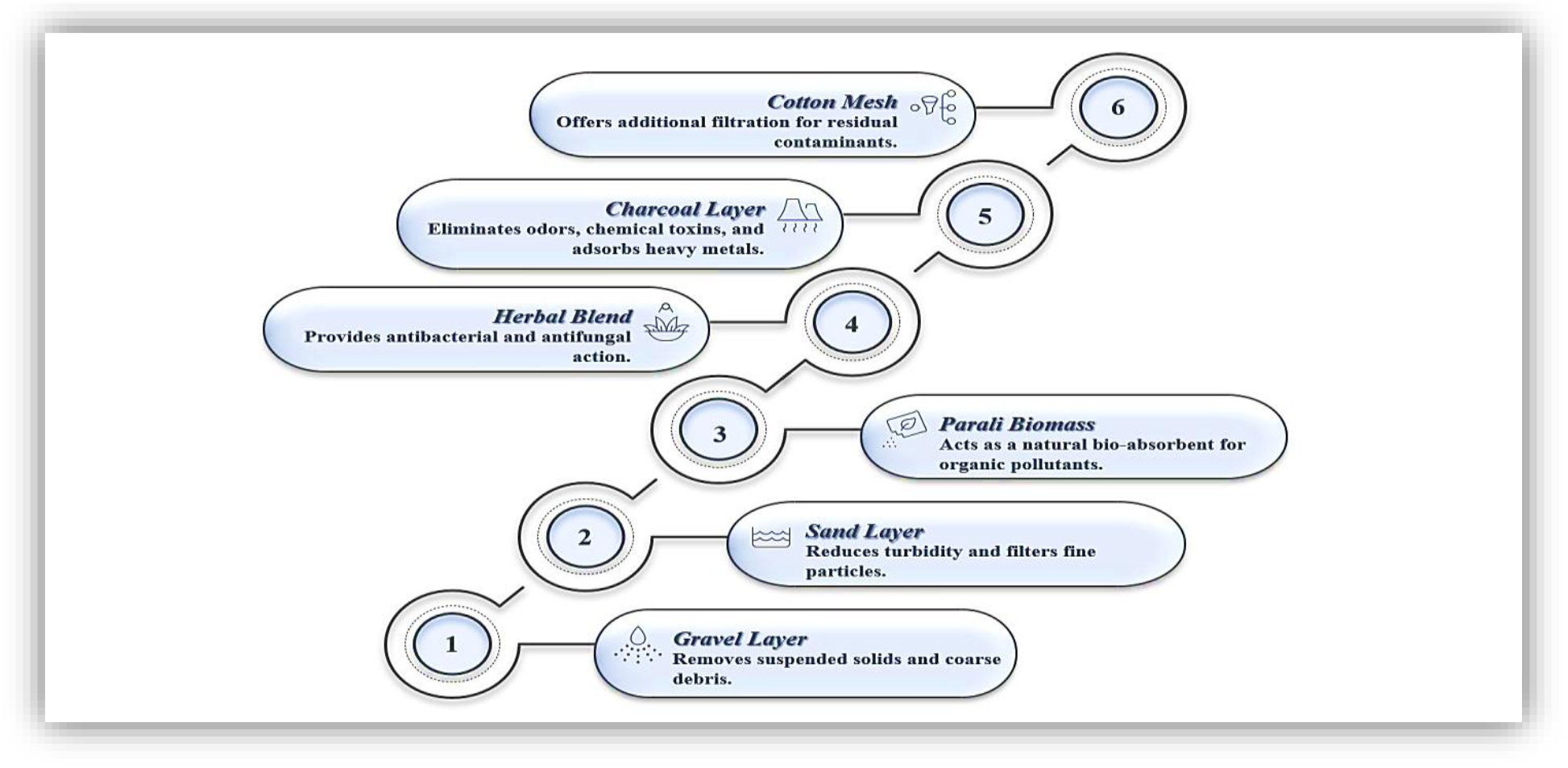
The six-layer herbal-biomass filtering system’s flowchart shows how physical, chemical, and microbiological impurities are removed one after the other.

**Figure 2:**
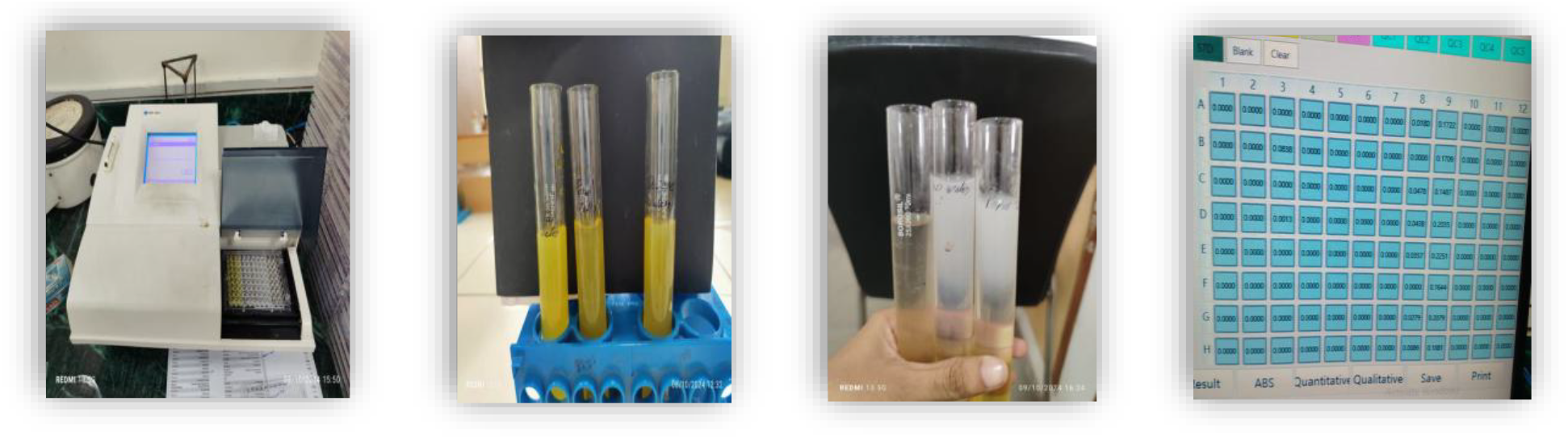
Using the Spectrophotometric Method to Perform the COD Test and Present the Results.

**Figure 3:**
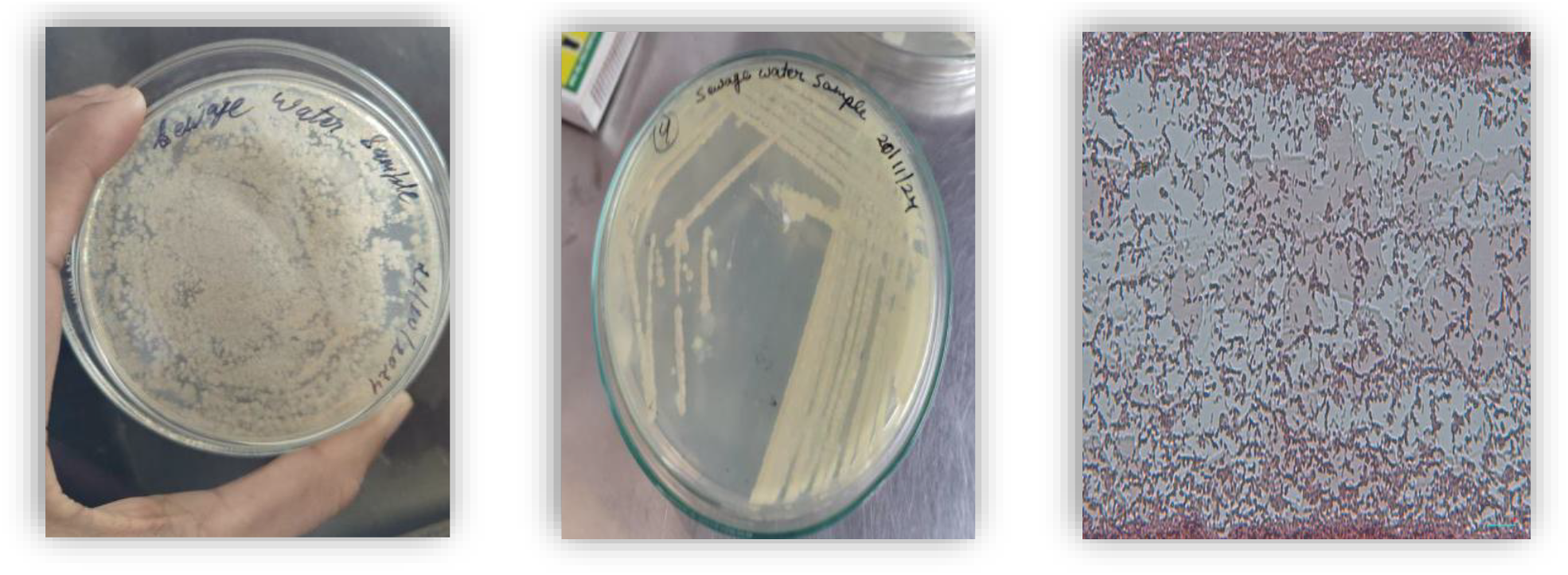
Gram-positive bacteria are revealed by microbial culture and Gramme staining findings.

**Figure 4:**
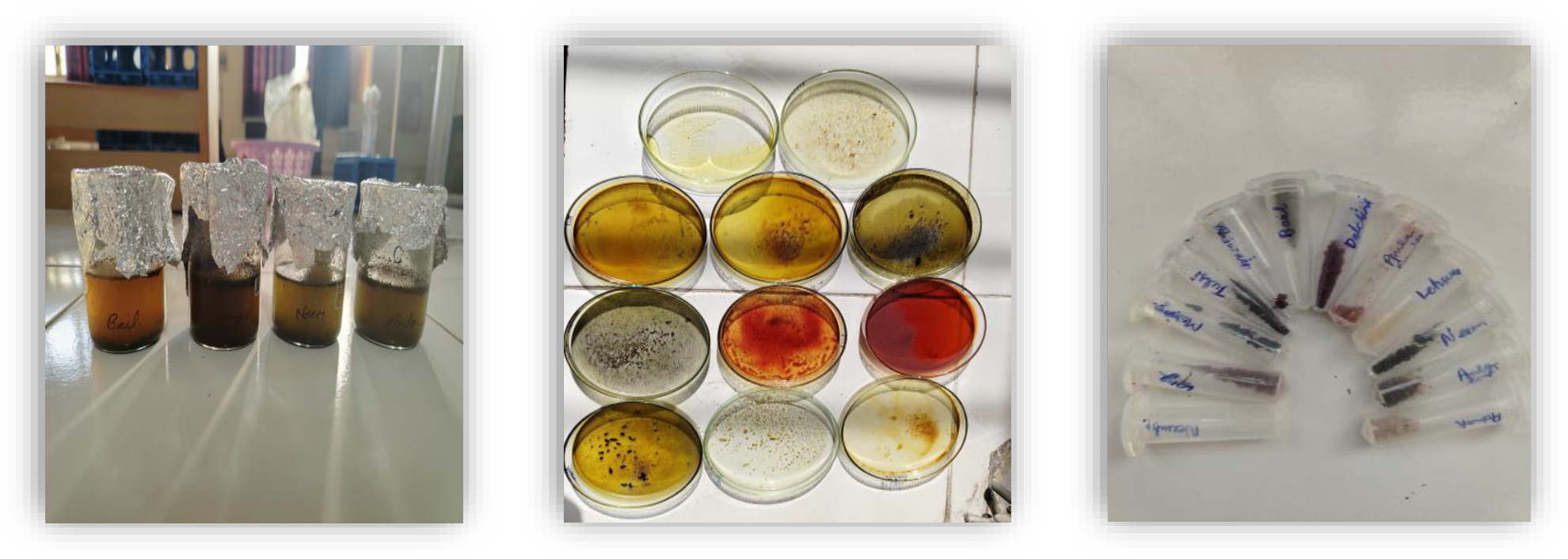
Prepared Herbal Extracts for Testing and Microbial Treatment.

**Figure 5:**
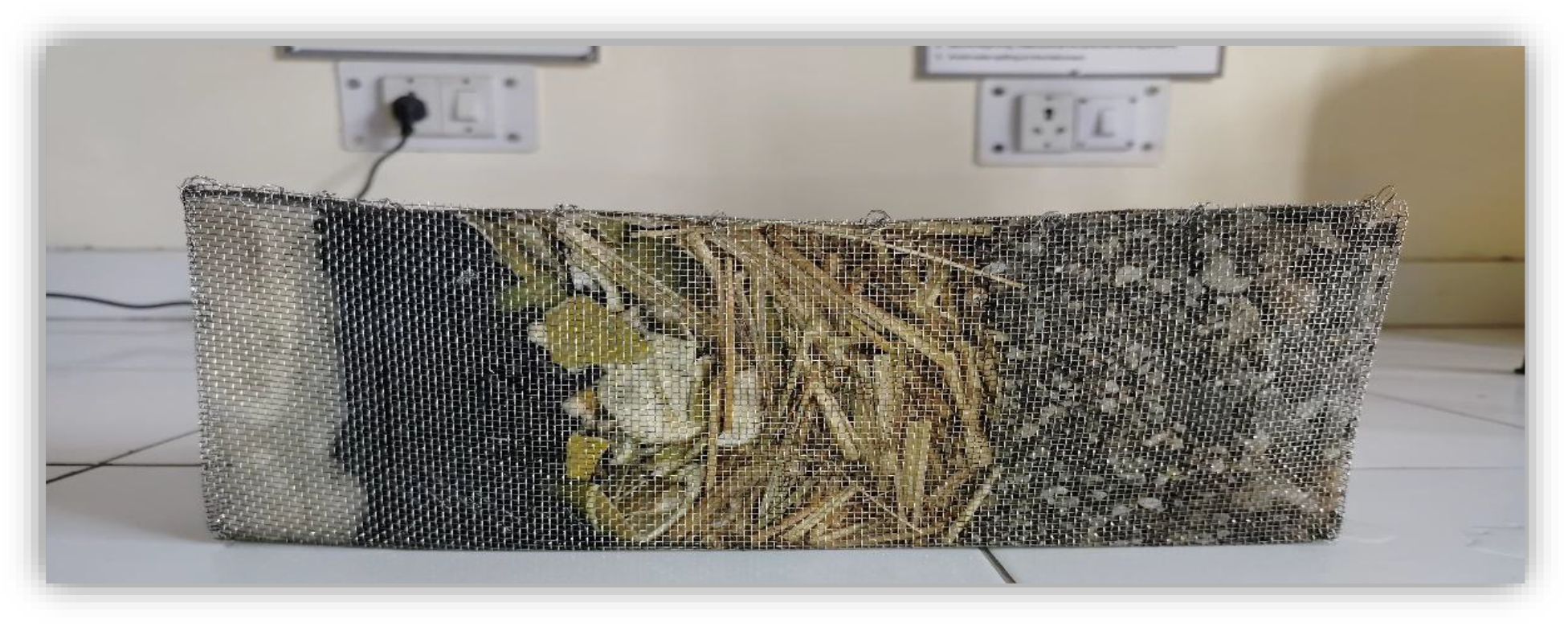
The filtering unit’s 1:4 scale tiny prototype was created in a lab.

**Figure 6:**
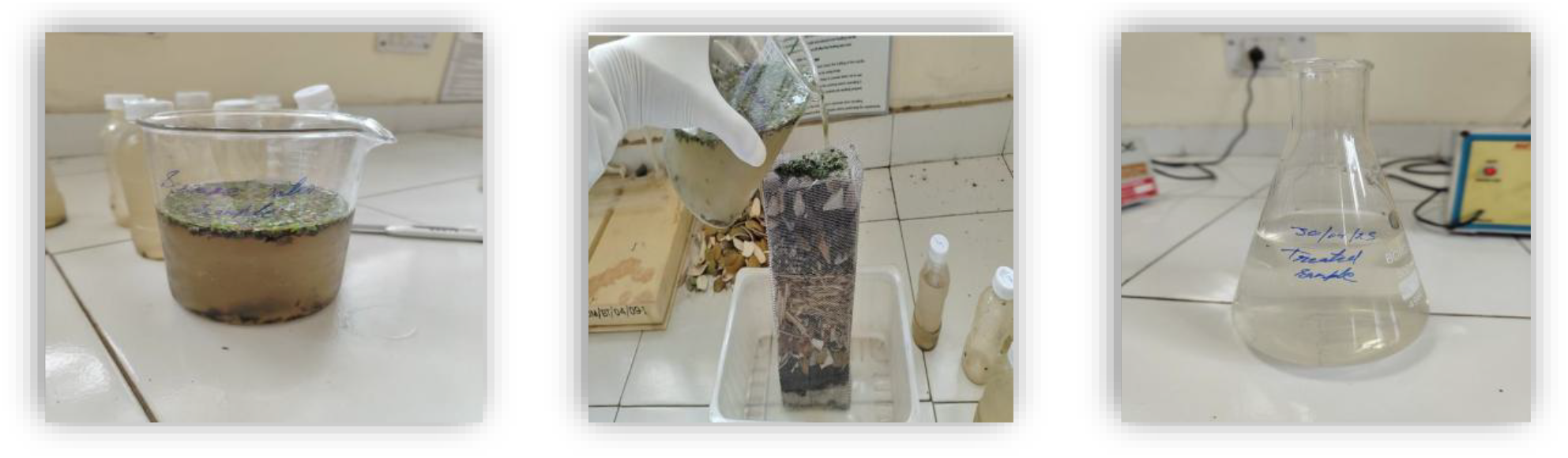
The miniature prototype created for water sample filtering performed successfully in the lab.

**Figure 7:**
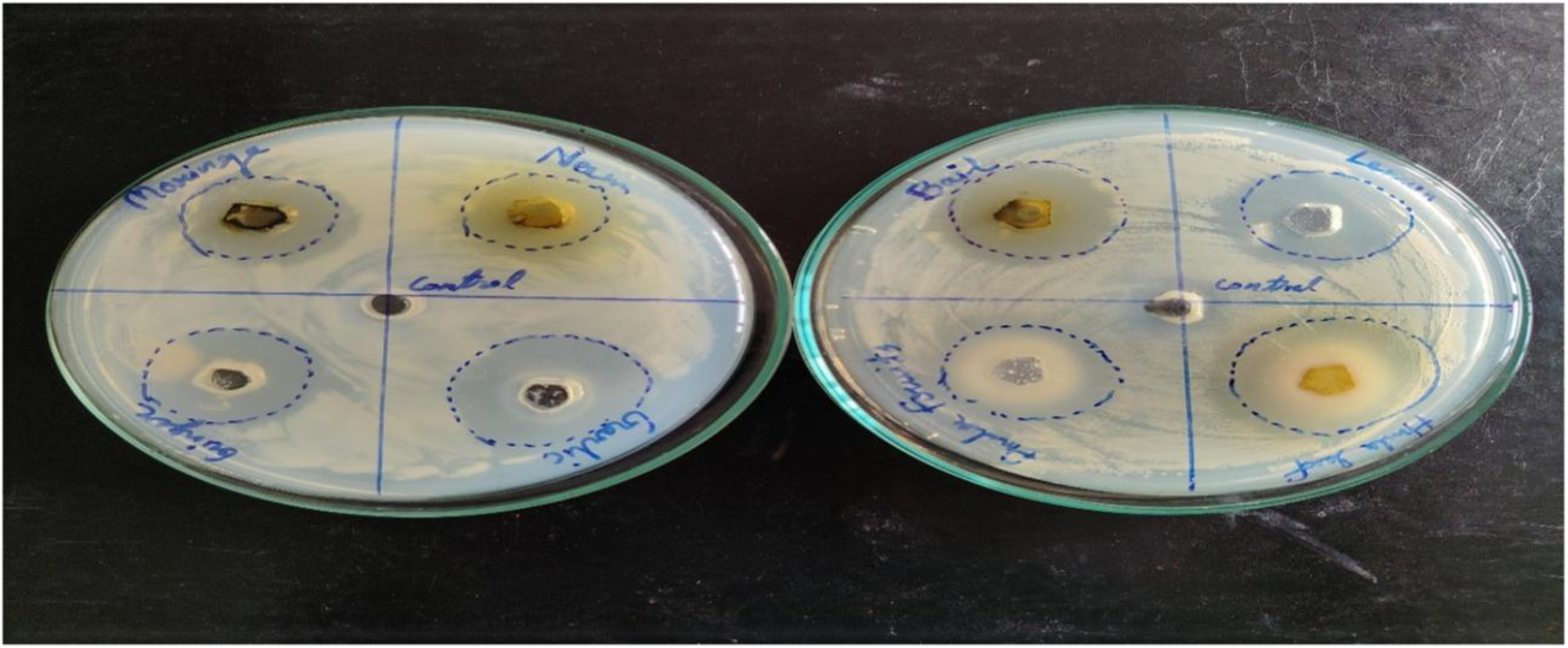
Neem, Moringa, Ginger, Garlic, Bael, Lemon, Basil, and Amla are herbal extracts that suppress the growth of multidrug-resistant (MDR) Streptobacillus moniliformis.

**Figure 8:**
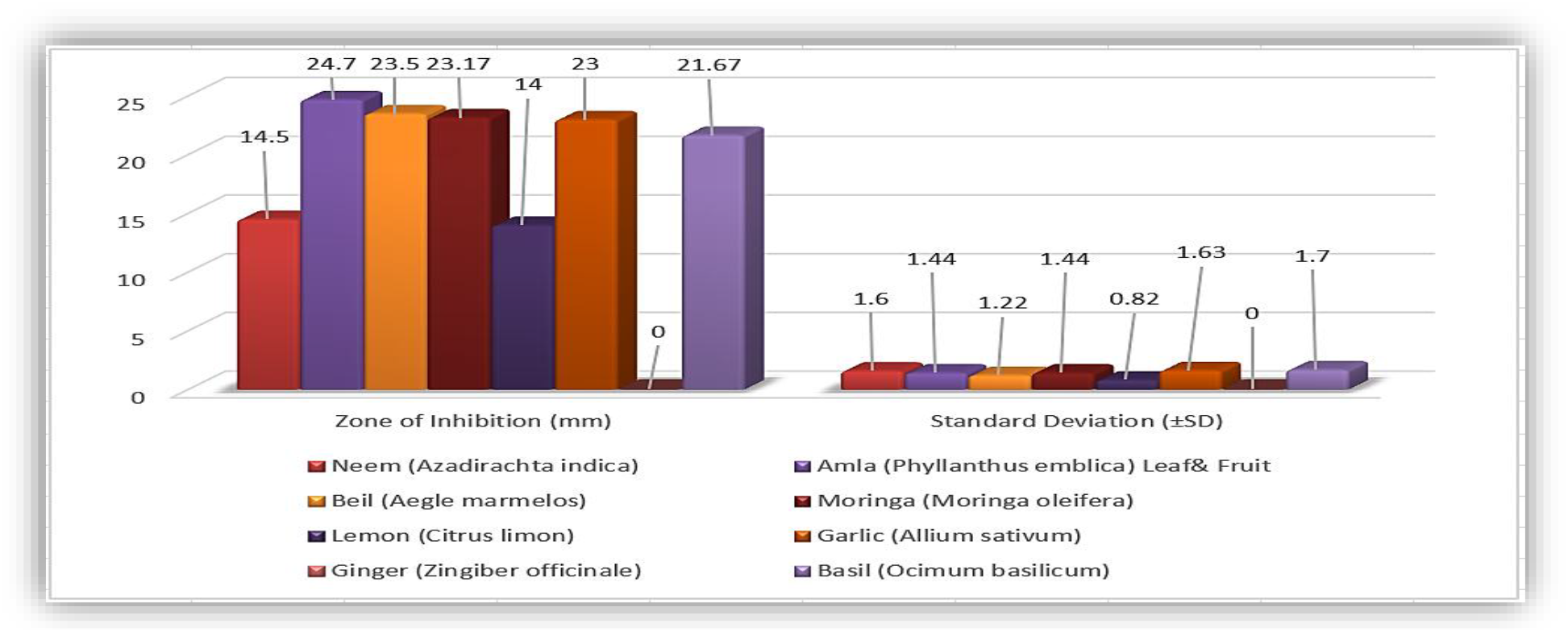
Assessment of herbal extracts’ antibacterial capabilities against Streptobacillus moniliformis.

### 3.3. Water Quality Improvement After Treatment

Water quality significantly improved after treatment with the six-layer prototype filter unit, producing results similar to membrane-based systems without requiring energy, expensive materials, or reject water production. The water became remarkably clean as the turbidity reduced sharply from 300 NTU to 0.8 NTU. While biological oxygen demand (BOD) dropped from 200 mg/L to 3.5 mg/L, indicating significant biological stabilisation, chemical oxygen demand (COD) dropped from 1000 mg/L to 9 mg/L, indicating almost total elimination of oxidisable organic waste.

### 3.4. Microbial and Heavy Metal Reduction Efficiency

Starting with microbial identification, a thorough experimental protocol was performed to evaluate the six-layer prototype filter’s effectiveness. Gramme staining was used to confirm the presence of Gram-positive bacteria in water samples, including tap and sewage water, after they were cultivated on nutrient agar. A microbial load of 2,000,000 CFU/mL in sewage water and 50 CFU/mL in tap water was found by colony-forming unit (CFU) enumeration. Select botanicals, including bael leaves, neem leaves, lemon peels, garlic, ginger, moringa, and amla, were soaked in distilled water for 24 and 48 hours in order to create herbal extracts for antibacterial testing. The agar well diffusion technique was then used to test these extracts against Streptobacillus moniliformis. The findings demonstrated that extracts steeped for 48 hours had much higher antibacterial activity.

Neem showed bactericidal effects, lemon peels and bael leaves created the biggest inhibition zones, while other botanicals showed moderate inhibition. After microbiological testing, untreated tap water with 0.02 mg/L of lead was used to assess heavy metal reduction.

Lead levels were lowered to undetectable levels after passing through the prototype. This was ascribed to the layers of cotton mesh and charcoal’s ability to adsorb heavy metals and chemical pollutants. The six-stage filtering system’s incorporation of herbal antimicrobials and bioabsorbent layers (parali biomass) produced total microbial and heavy metal removal, confirming the prototype’s promise as an affordable and environmentally friendly water purification method. Following treatment, the CFU counts in both cases dropped to 0 CFU/mL, indicating total microbial eradication (*Table 3*).

**Table 3:** Microbial Analysis Findings for a Sample of Treated and Untreated Water.

| *Sample Type* | *Untreated Load (CFU/mL)* | *Treated Load (CFU/mL)* |
| --- | --- | --- |
| Sewage Water | 2,000,000 | 0 (Negligible) |
| Tap Water | 50 | 0 (Negligible) |

### 3.5. Prototype Validation

The six-layer prototype filter (gravel → sand → parali biomass → herbal extracts → charcoal → cotton) showed encouraging results in water purification at the 1:4 scale, making it especially appropriate for use in rural areas. Microbial reduction was over 70% at intermediate filtering stages, and 0 CFU/mL at the final output indicated total microbial removal. In addition to efficiently adsorbing lead and lowering its concentration to undetectable levels, the charcoal-parali biomass combination also stabilised the pH between 7.66 and 6.9, bringing the water closer to neutrality. The system’s dependability was confirmed by the continuous removal of turbidity, chemical oxygen demand (COD), and biological oxygen demand (BOD) throughout several trials. This gravity-driven, community-operable design is more suited for decentralised rural environments than reverse osmosis or ultrafiltration systems, which need constant electricity and skilled maintenance. The concept is unusual since it addresses environmental and water quality issues by combining herbal antimicrobials with parali biomass, an agricultural leftover that is frequently burnt in open fields. ***Figure 10*** shows the sharp contrast between untreated and treated samples, whereas ***Figure 9*** compares pond water before and after treatment to show the improvement in clarity and decrease in microbial load. Because the system uses inexpensive, locally accessible materials, community involvement further guaranteed operability and acceptability while bolstering its scalability. Limitations still exist despite its success in lab settings, such as the absence of long-term durability data and full-scale deployment. Long-term field tests, multi-village implementation, and ratio optimisation for herbal extracts should be the main areas of future study. The multi-layer system provided a complete remedy for physical, chemical, and microbiological contamination; parali biomass improved adsorption capacity, while plant extracts formed a potent antibacterial barrier. All things considered, these results demonstrate that the herbal multi-barrier prototype is a practical, reasonably priced, and ecologically friendly substitute for traditional filtration technologies. It has the potential to greatly enhance public health and advance sustainable water management in rural areas.

**Figure 9:**
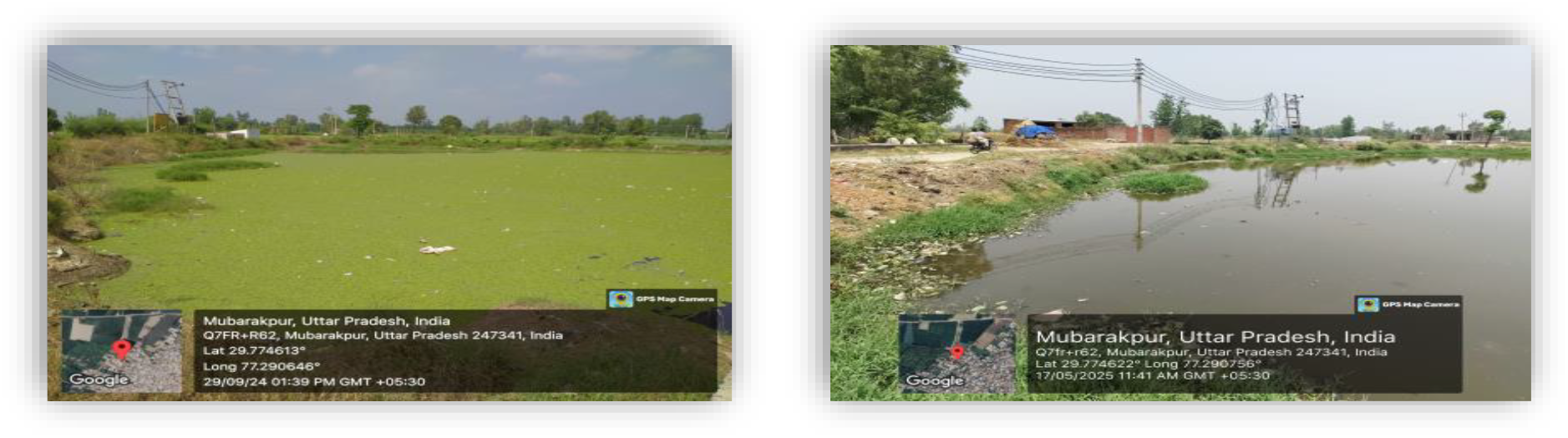
Using herbal antimicrobials and parali biomass, a prototype-based pond treatment demonstrated enhanced water quality and a decrease in microbial load.

**Figure 10:**
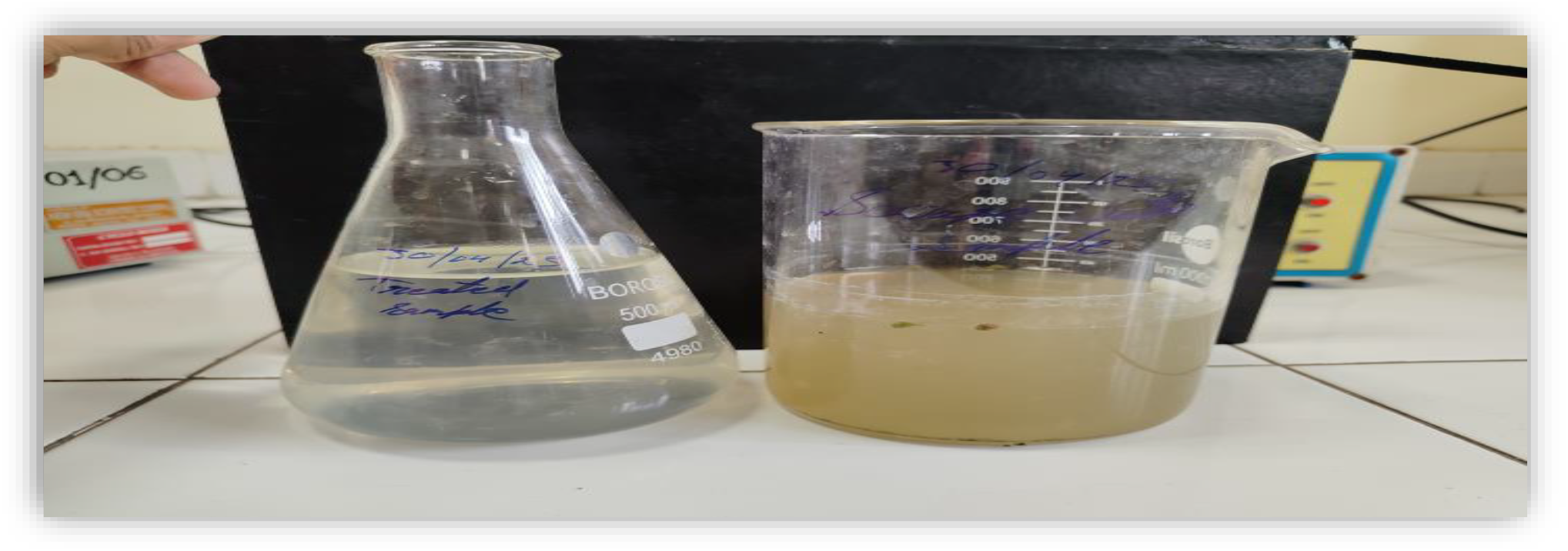
Comparison of treated (clear) and untreated (contaminated) water samples.

## Conclusion

This study, which was conducted in the hamlet of Mubarakpur, focusses on the development and assessment of an inexpensive, herbal-based water filtering system intended to reduce rural water pollution. Surveys, water sampling, and laboratory testing revealed significant contamination; untreated sewage had bacterial loads as high as 2,000,000 CFU/mL, turbidity of 300 NTU, COD of 1000 mg/L, and BOD of 200 mg/L. Even tap water was discovered to have lead levels, underscoring the pressing need for an affordable filtering system. According to herbal efficacy studies, neem extracts had potent bactericidal qualities, but bael leaves and lemon peels shown substantial antibacterial inhibition. In light of these findings, a six-layer filtering prototype was developed using charcoal, cotton fabric, parali biomass, gravel, sand, and herbal extracts. Laboratory validation showed remarkable improvements, including a reduction in turbidity to 0.8 NTU, a reduction in COD to 9 mg/L, a reduction in BOD to 3.5 mg/L, a reduction in bacterial load to 0 CFU/mL, and the total elimination of detectable lead. The system’s effectiveness stems from its innovative combination of herbal treatments and agricultural waste, which promotes environmental sustainability and rural health. Its gravity-driven, community-operable design ensures cost and ease of maintenance. Long-term performance, sensitivity to seasonal variations, and widespread acceptance are still problems, though. Future studies should focus on extensive field testing, optimising herbal formulations, and replication across many communities to verify scalability and durability. All things considered, the study demonstrates a practical, sustainable approach to water purification that also promotes environmental resilience and the management of agricultural residues in rural regions.

